# Analysis of the ARTIC version 3 and version 4 SARS-CoV-2 primers and their impact on the detection of the G142D amino acid substitution in the spike protein

**DOI:** 10.1101/2021.09.27.461949

**Authors:** James J. Davis, S. Wesley Long, Paul A. Christensen, Randall J. Olsen, Robert Olson, Maulik Shukla, Sishir Subedi, Rick Stevens, James M. Musser

**Affiliations:** Division of Data Science and Learning, Argonne National Laboratory, 9700 S. Cass Ave., Lemont, Illinois, 60439; University of Chicago Consortium for Advanced Science and Engineering, 5801 South Ellis Avenue, Chicago, Illinois, 60637; Center for Infectious Diseases, Laboratory of Molecular and Translational Human Infectious Diseases Research, Department of Pathology and Genomic Medicine, Houston Methodist Research Institute and Houston Methodist Hospital, 6565 Fannin Street, Houston, Texas, 77030; Departments of Pathology and Laboratory Medicine, and Microbiology and Immunology, Weill Cornell Medical College, 1300 York Avenue, New York, New York, 10065; Computing, Environment and Life Sciences Directorate, Argonne National Laboratory, Argonne, Illinois, USA, 60439; Department of Computer Science, University of Chicago, Chicago, Illinois, 60637

**Author notes:** Contributed equally to this work (James J. Davis, S. Wesley Long). Address correspondence to James J. Davis, Ph.D., Division of Data Science and Learning, Argonne National Laboratory, 9700 S. Cass Ave., Lemont, Illinois, 60439. **Disclosures:** None.

## Abstract

The ARTIC Network provides a common resource of PCR primer sequences and recommendations for amplifying SARS-CoV-2 genomes. The initial tiling strategy was developed with the reference genome Wuhan-01, and subsequent iterations have addressed areas of low amplification and sequence drop out. Recently, a new version (V4) was released, based on new variant genome sequences, in response to the realization that some V3 primers were located in regions with key mutations. Herein, we compare the performance of the ARTIC V3 and V4 primer sets with a matched set of 663 SARS-CoV-2 clinical samples sequenced with an Illumina NovaSeq 6000 instrument. We observe general improvements in sequencing depth and quality, and improved resolution of the SNP causing the D950N variation in the spike protein. Importantly, we also find nearly universal presence of spike protein substitution G142D in Delta-lineage samples. Due to the prior release and widespread use of the ARTIC V3 primers during the initial surge of the Delta variant, it is likely that the G142D amino acid substitution is substantially underrepresented among early Delta variant genomes deposited in public repositories. In addition to the improved performance of the ARTIC V4 primer set, this study also illustrates the importance of the primer scheme in downstream analyses.

**Importance:** ARTIC Network primers are commonly used by laboratories worldwide to amplify and sequence SARS-CoV-2 present in clinical samples. As new variants have evolved and spread, it was found that the V3 primer set poorly amplified several key mutations. In this report, we compare the results of sequencing a matched set of samples with the V3 and V4 primer sets. We find that adoption of the ARTIC V4 primer set is critical for accurate sequencing of the SARS-CoV-2 spike region. The absence of metadata describing the primer scheme used will negatively impact the downstream use of publicly available SARS-Cov-2 sequencing reads and assembled genomes.

## Observation

The ARTIC Network is a consortium dedicated to providing tools to support global viral epidemiology efforts through low cost genomic sequencing (https://artic.network). Early in the SARS-CoV-2 pandemic, the consortium developed a set of primers (ARTIC V1) designed to completely sequence the SARS-CoV-2 genome with overlapping 400-bp amplicons. Shortcomings identified in the V1 protocol, primarily due to regions of amplicon drop out, led to two more iterations of primer design, which resulted in ARTIC V3 being used by many laboratories in 2020 and into 2021 (1). As the COVID-19 pandemic continued, new variants emerged with unique mutations and enhanced transmissibility (2–4). Some of these mutations occurred in primer binding sites in genes that encode key proteins such as spike protein, resulting in amplicon dropout and poor sequence coverage in critical regions. ARTIC V4 was a new set of tiling primers posted on June 18, 2021 (https://github.com/artic-network/artic-ncov2019/tree/master/primer_schemes/nCoV-2019), designed using multiple variant sequences as input to address these issues. In particular, there were spike protein amino acid changes common to the Beta, Delta, and Gamma variants that occurred in known V3 primer binding sites, including G142D (Delta) in the 2_Right primer, the 241/243del (Beta) that occurs in the 74_Left primer, and the K417N (Beta) or K417T (Gamma) which occur in the 76_Left primer (https://community.artic.network/t/sars-cov-2-version-4-scheme-release/312).

From the beginning of the pandemic, we strived to sequence all patient samples with SARS-CoV-2 in the Houston Methodist Hospital system, a large 2,500-bed healthcare system in Houston, Texas, USA. In the summer of 2021, we experienced a massive surge of patients with COVID-19 that corresponded with an increase in Delta variant cases (4). Although we have been using the ARTIC primer sets throughout the pandemic, we elected to validate the ARTIC V4 primer set prior to adopting the new protocol. We chose a random set of 663 SARS-CoV-2 clinical samples isolated between July 2-18, 2021, and each of the 663 samples was amplified using the V3 and V4 primers. SARS-CoV-2 nucleic acid present in the samples was amplified by methods described previously (5, 6). Samples were sequenced with an Illumina NovaSeq 6000 instrument. Paired-end reads for both the V3 and V4 amplified samples were assembled with the assembly service of the National Institute of Allergy and Infectious Diseases (NIAID)-funded Bacterial and Viral Bioinformatics Resource Center (BV-BRC) (https://www.bv-brc.org), which follows the One-Codex workflow (https://github.com/onecodex/sars-cov-2). The workflow uses seqtk version 1.3-r116 for quality trimming (https://github.com/lh3/seqtk.git); minimap version 2.143 for aligning reads against Wuhan-Hu-1 (NC_045512.2) (7); samtools version 1.11 for sequence and file manipulation(8); and iVar version 1.2.2(9) for primer trimming and SNP calling. Default parameters were used in all cases except that the maximum read depth in mpileup was limited to 8,000, and the minimum read depth for a variant call in iVar was set to 3. Lineages were assigned with Pangolin version 3.1.11 using pangoLearn module 2021-08-24 (https://cov-lineages.org/resources/pangolin.html)(10). All sequencing reads are available at SRA under bioproject PRJNA767338.

Overall, we observed considerable improvement in sequence quality of the V4 assemblies relative to V3. The median read depths tended to be higher at each nucleotide position **(Fig 1A)** and at each primer position **(Figs 1B and 1C).** Notably, the V3 region of low coverage spanning approximately nucleotide positions 22,320–22,530 (corresponding to V3 primer pair 74) located in the spike gene, is corrected in V4. Consistent with previous analysis of ARTIC V4 (https://community.artic.network/t/sars-cov-2-version-4-scheme-release/312),we observe an area of slightly lower coverage in the V4 sequences at approximate nucleotide positions 26,950–27,180 (corresponding to V4 primer 90), but this was less problematic because we observed fewer assemblies with runs of ambiguous base calls in the V4 set.

**FIG 1.**
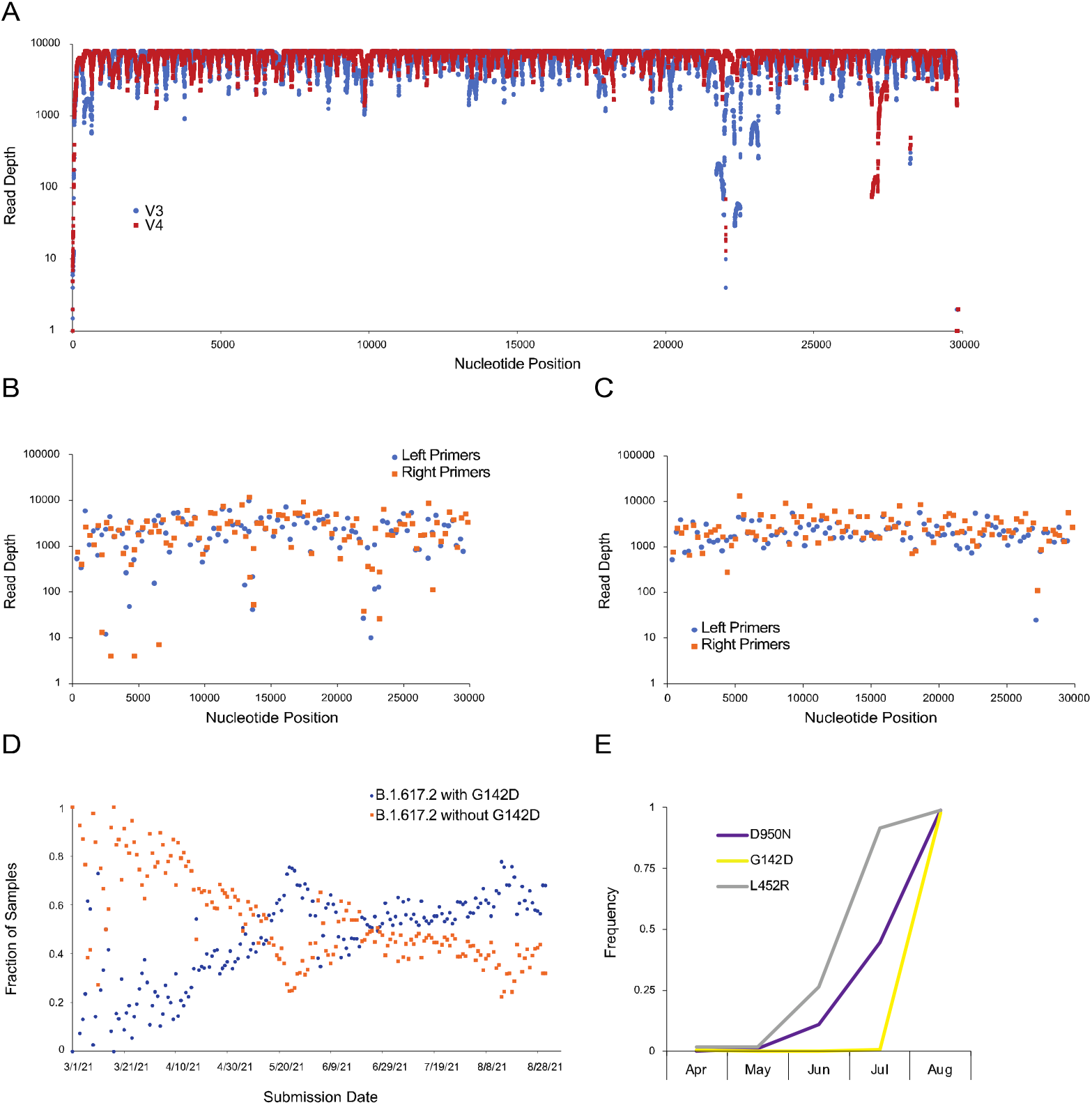
Sequencing artifact analysis of spike protein amino acid position 142. (A) Median read depths at each nucleotide position for the assembly of the set of 663 V3 (blue) and V4 (red) samples. (B) Median read depth at each primer for the set of 663 V3 assemblies. (C) Median read depth at each primer for the set of 663 V4 assemblies. Orange squares are right primers and blue circles are left primers. (D) The fraction of B.1.617.2 sequences in GISAID with (blue) and without (orange) G142D through August 31, 2021. (E) Frequency of L452R (gray), D950N (blue), and G142D (orange) amino acid substitutions observed in SARS-CoV-2-positive samples from April through August of 2021. L452R, D950N, and G142D are hallmark amino acid substitutions of the Delta variant.

Among the 663 samples, 53 had different pangolin calls in the V3 versus V4 assemblies **(Table 1**). None of these 53 sample pairs had identical spike protein sequences. The most common nucleotide difference occurs at nucleotide position 21,987, which is the G to A transition that causes the G142D amino acid spike variant. The second most common SNP occurs in position 24,410, which causes the D950N amino acid variant. Three hundred sixty-eight of the V3 assemblies had an ambiguous base at this position compared with only 3 of the V4 assemblies. Except for the ends of the assembled sequences which can be jagged, and therefore ambiguous, 3 samples (MCoV-49081, MCoV-50268 and MCoV-49000) differed only at position 21,987 (G142D) SNP in the V3 versus V4 assemblies. These three V3 assemblies were classified by pangolin as being either AY.15 or AY.24. When sequenced by V4, they were all classified as B.1.617.2. A fourth sample (MCoV-50188) differed only at positions 21,987 (G142D) and 24,410 (D950N). This also changed the pangolin classification from AY.15 in V3 to B.1.617.2 in V4. The remaining 53 samples with differing pangolin lineages had 3 or more SNPs in the V3 versus V4 run. In this set of 663 genomes, none of the V4 assemblies with a complete spike protein that are classified as being a Delta or a Delta sublineage have the ancestral glycine at position 142.

**Table 1.** Distribution of Pangolin lineages in the set of 663 samples that were sequenced using either the ARTIC version 3 or version 4 primers.

| **Lineage** | **ARTIC Version 3** | **ARTIC Version 4** |
| --- | --- | --- |
| AY.10 | 2 | 1 |
| AY.12 | 4 | 0 |
| AY.13 | 2 | 2 |
| AY.14 | 1 | 1 |
| AY.15 | 5 | 0 |
| AY.2 | 4 | 7 |
| AY.20 | 3 | 3 |
| AY.21 | 1 | 0 |
| AY.24 | 5 | 1 |
| AY.25 | 124 | 126 |
| AY.3 | 40 | 36 |
| AY.3.1 | 6 | 7 |
| AY.4 | 4 | 0 |
| B.1 | 0 | 1 |
| B.1.1 | 0 | 1 |
| B.1.1.7 | 40 | 40 |
| B.1.575 | 1 | 0 |
| B.1.617.2 | 375 | 398 |
| B.1.621 | 4 | 4 |
| B.1.621.1 | 1 | 1 |
| B.1.625 | 0 | 1 |
| B.1.628 | 11 | 11 |
| B.1.637 | 1 | 1 |
| C.37 | 4 | 3 |
| P.1 | 9 | 9 |
| P.1.10 | 1 | 1 |
| None | 15 | 8 |

Public repositories such as GISAID (11) and the INSDC resources (12) host SARS-CoV-2 genome sequences collected globally. These databases have been indispensable for epidemiological analyses, early identification of variants of concern, and downstream translational research activities such as vaccine formulation. From June 2021 through August 2021, the rapid increase in the G142D amino acid substitution present in Delta variants in public repositories appeared to indicate a rapid evolutionary sweep **(Fig 1D, Table S1)** bearing resemblance to previous evolutionary sweeps, including the D614G substitution in 2020(13), B.1.1.7 (Alpha) last fall and winter (2, 5), and Delta this spring and summer (3, 4). However, our data lead us to conclude that the sharp uptick in spike protein G142D was caused by community adoption of the V4 primers. Indeed, when we examine 12,441 samples from Houston Methodist patients collected since April of 2021, comparing the occurrence of G142D with L452R (another hallmark Delta substitution in spike), it becomes clear that the G142D uptick is an artifact that corresponds precisely with our adoption of the V4 primers in mid-July 2021 **(Fig 1E).** Indeed, only 2 Delta variant genomes collected after July 1, 2021 had the ancestral glycine at position 142 (4).

## Conclusion

The results of this study are consistent with those published by the ARTIC network (https://community.artic.network/t/sars-cov-2-version-4-scheme-release/312). We observe substantially improved sequence quality, including higher median read depths and fewer regions of ambiguous base calls in the V4 assemblies compared with V3. We also observe that the ancestral glycine at spike position 142 is extremely rare in Delta variants collected in Houston. This study indicates that the primer scheme used for amplifying and sequencing SARS-CoV-2 genomes is an important consideration for interpreting epidemiological data and identifying variants of concern.

## Acknowledgments

We thank Liliana Brown, Elodie Ghedin, Wiriya Rutvisuttinunt, and many other colleagues for persistently encouraging this project. We thank Emily Dietrich and Heather McConnell for help with manuscript and figure preparation. We thank our dedicated SARS-CoV-2 genomic sequencing team, including Akanksha Batajoo, Jessica Cambric, Ryan Gadd, Regan Mangham, Matthew Ojeda Saavedra, Sindy Pena, Layne Pruitt, Kristina Reppond, Madison N. Shyer, Rashi M. Thakur, Trina Trinh, and Prasanti Yerramilli for their tireless efforts throughout the pandemic in generating the sequencing data. This work was supported by the Houston Methodist Academic Institute Infectious Diseases Fund and many generous Houston philanthropists. JJD, RO, and MS were funded in whole or in part with Federal funds from the National Institute of Allergy and Infectious Diseases, National Institutes of Health, Department of Health and Human Services, under Contract No. 75N93019C00076 to principal investigator Rick Stevens.

## References

1. Tyson JR, James P, Stoddart D, Sparks N, Wickenhagen A, Hall G, Choi JH, Lapointe H, Kamelian K, Smith AD, Prystajecky N, Goodfellow I, Wilson SJ, Harrigan R, Snutch TP, Loman NJ, Quick J. 2020. Improvements to the ARTIC multiplex PCR method for SARS-CoV-2 genome sequencing using nanopore. bioRxiv doi:10.1101/2020.09.04.283077:2020.09.04.283077.

2. Davies NG, Abbott S, Barnard RC, Jarvis CI, Kucharski AJ, Munday JD, Pearson CA, Russell TW, Tully DC, Washburne AD. 2021. Estimated transmissibility and impact of SARS-CoV-2 lineage B. 1.1. 7 in England. Science 372.

3. Petra M, Steven K, Mahesh Shanker D, Guido P, Bo M, Swapnil M, Charlie W, Thomas M, Isabella F, Rawlings D, Dami AC, Sujeet S, Rajesh P, Robin M, Meena D, Shantanu S, Kalaiarasan P, Radhakrishnan VS, Adam A, Niluka G, Jonathan B, Oscar C, Partha C, Priti D, Daniela C, Tom P, Dr Chand W, Neeraj G, Raju V, Meenakshi A, The Indian S-C-GC, Citiid-Nihr BioResource Covid-19 Collaboration AM, o Hyeon L, Wendy SB, Samir B, Seth F, Leo J, Partha R, Anurag A, Ravindra KG. 2021. Nature Portfolio doi:10.21203/rs.3.rs-637724/v1.

4. Christensen PA, Olsen RJ, Long SW, Subedi S, Davis JJ, Hodjat P, Walley DR, Kinskey JC, Saavedra MO, Pruitt L, Reppond K, Shyer MN, Cambric J, Gadd R, Thakur RM, Batajoo A, Mangham R, Pena S, Trinh T, Yerramilli P, Nguyen M, Olson R, Snehal R, Gollihar J, Musser JM. 2021. Delta variants of SARS-CoV-2 cause significantly increased vaccine breakthrough COVID-19 cases in Houston, Texas. medRxiv doi:10.1101/2021.07.19.21260808:2021.07.19.21260808.

5. Long SW, Olsen RJ, Christensen PA, Subedi S, Olson R, Davis JJ, Saavedra MO, Yerramilli P, Pruitt L, Reppond K. 2021. Sequence Analysis of 20,453 Severe Acute Respiratory Syndrome Coronavirus 2 Genomes from the Houston Metropolitan Area Identifies the Emergence and Widespread Distribution of Multiple Isolates of All Major Variants of Concern. The American Journal of Pathology.

6. Olsen RJ, Christensen PA, Long SW, Subedi S, Hodjat P, Olson R, Nguyen M, Davis JJ, Yerramilli P, Saavedra MO. 2021. Trajectory of Growth of Severe Acute Respiratory Syndrome Coronavirus 2 (SARS-CoV-2) Variants in Houston, Texas, January through May 2021, Based on 12,476 Genome Sequences. The American Journal of Pathology.

7. Li H. 2018. Minimap2: pairwise alignment for nucleotide sequences. Bioinformatics 34:3094–3100.

8. Li H, Handsaker B, Wysoker A, Fennell T, Ruan J, Homer N, Marth G, Abecasis G, Durbin R. 2009. The sequence alignment/map format and SAMtools. Bioinformatics 25:2078–2079.

9. Grubaugh ND, Gangavarapu K, Quick J, Matteson NL, De Jesus JG, Main BJ, Tan AL, Paul LM, Brackney DE, Grewal S. 2019. An amplicon-based sequencing framework for accurately measuring intrahost virus diversity using PrimalSeq and iVar. Genome biology 20:1–19.

10. Rambaut A, Holmes EC, O’Toole Á, Hill V, McCrone JT, Ruis C, du Plessis L, Pybus OG. 2020. A dynamic nomenclature proposal for SARS-CoV-2 lineages to assist genomic epidemiology. Nature microbiology 5:1403–1407.

11. Shu Y, McCauley J. 2017. GISAID: Global initiative on sharing all influenza data–from vision to reality. Eurosurveillance 22:30494.

12. Arita M, Karsch-Mizrachi I, Cochrane G. 2021. The international nucleotide sequence database collaboration. Nucleic Acids Research 49:D121–D124.

13. Korber B, Fischer WM, Gnanakaran S, Yoon H, Theiler J, Abfalterer W, Hengartner N, Giorgi EE, Bhattacharya T, Foley B. 2020. Tracking changes in SARS-CoV-2 spike: evidence that D614G increases infectivity of the COVID-19 virus. Cell 182:812–827. e19.

